# Mutualistic relationships between male inflorescences of *Elaeis guineensis* Jacquin and its pollinator *Elaeidobius kamerunicus* Faust (Coleoptera: Curculionidae: Derelominae) Comparison between two cultivars for *E. kamerunicus* progeny emergence in North Sumatra Indonesia

**DOI:** 10.64898/2026.10.14.759277

**Authors:** L. Beaudoin-Ollivier, F. Jacob, C. Madec, D. Afandi, I. Syahputra, H. Rey, Y. Dumont

## Abstract

*Elaeidobius kamerunicus* completes its life cycle on male inflorescences while ensuring pollen transfer to female inflorescences. Over the past decades, oil palm breeding has prioritized yield potential over male inflorescence production. In young plantations, scare male inflorescences may result in insufficient pollen and breeding sites for *E. kamerunicus*, reducing or eliminating local populations. Consequently, reduced pollination lowers fruit set and yield. Our study assessed weevil emergence phenology and potential from 74 post-anthesis male inflorescences of two cultivars: 31 from a super-male and 43 from a high-yielding experimental cultivar. Male spikelet morphology was compared using field and laboratory data collected in North Sumatra in Indonesia (2020-2022). Super-male palms had more spikelets (283 vs.174), greater total spikelet length (39 vs. 27 m) and higher *E. kamerunicus* emergence (17,500 vs. 13,100). The kinetic of emergence reveals that *E. kamerunicus* females emerged in a large number during 10 days and earlier than males which emerged during a longer period of time (18 days), indicative of protogyny in adult emergence. The predicted emergence kinetics indicate that *E. kamerunicus* from Experimental cultivar emerged earlier (1-2 days) than *E. kamerunicus* from Supermale and it also shows a difference in the timing of emergence between the two sexes. Thus, sex and cultivar jointly influence the emergence dynamics of *E. kamerunicus*. There was good evidence that the total length of the spikelets in an inflorescence positively influences the number of *E. kamerunicus* emergence events. We will discuss our results from the point of view of pollination.

## Introduction

The oil palm *Elaeis guineensis* Jacquin (Arecaes: Arecaceae) is a monoecious plant with allogamous reproduction that requires cross-pollination. The same individual produces both male and female inflorescences in alternating cycles of varying duration depending on genetic and environmental conditions (Forero et al. 2012). The pollination is both anemophilous or entomophilous (Hartley 1967; Sparnaaij 1969; Syed 1979, 1982) with the latter being considered of more importance for palm cultivation. *E. guineensis* pollination is an essential ecosystem service to ensure production of bunches in commercial plantations and is one of the main factors affecting production yield. Oil palm pollination mainly depends on *Elaeidobius kamerunicus* Faust 1878 (Coleoptera: Curculionidae: Derelominae), the main pollinator insect of oil palm in Indonesia since its successful introduction in the 1980’s (Whahid et al. 1987; Dhileepan 1994). The insects carrying pollens on their body travel out from the male inflorescences and visit receptive female inflorescence in anthesis on other palms drawn by the attractivity of volatile organic compounds. Its results a significant increase in fruit set and other bunch characters (Hutauruk et al. 1984; Susanto et al. 2007; Purba et al. 2010; Gintoron et al. 2023).

Development rate of immature stages of *E. kameruncius* (*EK*) has been described by Syed (1982). Newly emerged *EK* mate two to three days after emergence. The females lay their eggs in the male flowers, of which there are about a thousand per spikelet, and hence 2 to 300 thousand per inflorescence (Mariau et al. 1991). Oviposition starts within 2 days in the anthesising male spikelets. The oviposition period ranged 7 to 12 days (Dhileepan 1994). The eggs are deposited in the feeding pits by the female (Syed 1982). Egg lasts 1-2 days. There are three larval instars. After hatching the 1^st^ instar larval feeds on the dying tissues surrounding the oviposition site for 1-2 days. After it moults, the 2^nd^ instar moves to the base of the flower and feeds on the soft tissue for 1-2 days. It moults again and the 3^rd^ instar continue feeding before cutting a hole through the hard perianth near the base of the adjacent flower (5-9 days including the prepupal period). The 3^rd^ instar may consume 5 to 6 flowers (Syed 1982; Ponnamma et al. 2006). Pupation takes place in the consumed flower and it lasts for three days (Syed et al. 1982). The adults emerge 10-13 days later (Dhileepan 1994). The full development for the male is about 10-14 days and 9-14 days for the female (Syed 1979). Male average longevity is 16 days (7-24 days), while that of females was 11.7 days (5-21 days) (Dhileepan 1994). The weevil measures ± 4 mm long and ± 1.5 mm wide and is blackish brown in color (Syed 1982). There is a sexual dimorphism: the female has a smaller body size (2-3 mm), long rostrum and fine hairs. The male is longer (3-4 mm), shorter rostrum, more fine hairs on the edges of the abdomen and a protrusion at the base of the elytra (Fig. 1) (Latip et al. 2019).

**Figure 1.**
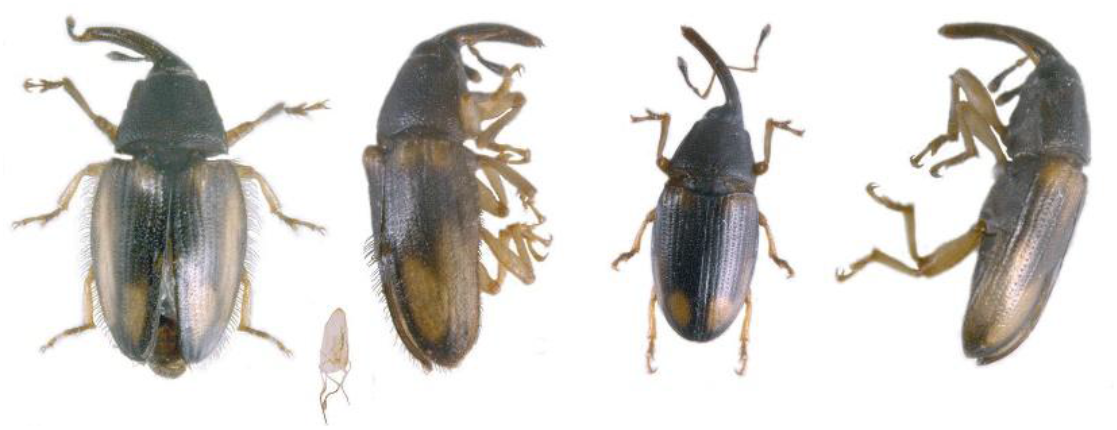
E. kamerunicus sex determination (from left to the right: EK male dorsal and profile view; EK female dorsal and profile view)

There has been concern, however, regarding the periodic occurrence of poor pollination and yield loss in certain Indonesian oil palm plantations. In commercial planting, the insect population decreased resulting in oil palm bunches with a very low fruit set value (Prasetyo & Susanto 2012). This is observed, especially in young oil palms that produce insufficient male inflorescence. Breeding program with new genotypes has successfully produced planting materials of high yielding materials in favor of female inflorescences. A reduction in the number of male inflorescences per area may have a drastically negative effect on the *EK* population. Pollination activity and yield may be influenced by the abundance of the *EK* population (Donough et al. 1996; Wahid & Kamarundin 1997; Prasetyo & Susanto 2019; Prasetyo et al. 2014). Factors that led to the problem were reported by Li et al. (2019).

The *E. guineensis* male inflorescence in anthesis is the main host for *EK* oviposition (Adaigbe et al. 2011). It represents a source of food, breeding sites and shelter for *EK* (Sambathkumar & Ranjith 2011; Syed 1979; Corley & Tinker 2008).

According to Beaudoin-Ollivier et al. (2026) the first estimates of 10 % of the immature stages occupying male inflorescences are interesting in light of the future estimation of *EK* progeny measured from the spikelets of experimental cultivars (EXP palms) compared to those from super mâle cultivars (SM palms) inflorescences. Thus, the relationship between the male inflorescence and the weevil population needs to be investigated to determine the potential of post-anthesis male inflorescence (PAMI) to produce *EK* imagos responsible of the pollination. One factors to ensure the sustainability of *EK* in the field is to measure the number of progenies that successfully emerge from the PAMI.

Weevil population has been estimated in several different ways (Chiu et al. 1986; Corley & Tinker 2016). Many authors estimated the weevil population visiting male inflorescence in anthesis (MIA) by collecting samples of spikelets from PAMI containing larvae and pupae, the spikelets being stored until adult weevils emerge and can be counted (Corley & Tinker 2016). The method commonly used to estimate progeny emergence is the stratified sampling of few spikelets of the MIA (Dhileepan 1994; Siswanto & Soetopo 2020; Mohamad et al. 2022) or the middle portion of the PAMI (Ponnamma et al. 2006).

Information regarding the phenology of *EK* emergence is limited to few spikelets per inflorescence in the literature. The present study aims to quantify the progenies emergence of *EK* emerging from entire PAMI and the factors that influence it such as the planting materials (EXP and SM cultivars), the morphology of the inflorescence, the time of sampling (year, month) and the climatic conditions for a period of three years.

## Material and methods

The present study was carried out for a period of two years from January 2020 to December 2022. The research was conducted on oil palm plantation at Bangun Bandar estate 2 (3°17’°03.12 ‘’N 99°01’57.73’’E) at the PT SOCFINDO Seeds Production Laboratory (SSPL) located at about 80 km from Medan in North Sumatra Province on Sumatra Is. in Indonesia. North Sumatra is characterized by two main seasons: a rainy season (October to April) and a dry season (May to September). The climate of the study site is hot humid tropical. The average temperature was 26°C, with a minimum of 21°C, a maximum of 36°C and about 80% relative humidity (HR) measured with an automatic Watch Dog 2900ET coupled with Spec9pro software (SDEC, France). Annual rainfall was around 2,500 mm, measured daily with a standard manual rain gauge.

The planting materials used in this study consisted of two *E. guineensis* cultivars: the “super male” (SM) cultivar clone SM-1 selected for its very high rate of male inflorescences; and an experimental cultivar (EXP) selected for its high rate of female inflorescences and high oil yield. Both cultivars were planted in a trial of 64 ha in May 2014. The objective of the trial was to improve pollination of the very female-biased EXP cultivar based on additional “super male” (SM) palms able to produce mainly male inflorescences. These SM palms were planted either in single (SD) or in double density (DD). For further description of the trial see Dumont et al. 2026. The current study was carried out on oil palms at 6 to 8-years after planting considering the two-planting materials SM and EXP. No chemicals were applied to the inflorescences during the study period.

### Experimental design and data collection

Systematic survey was employed to locate pre-anthesis male inflorescence on SM-SD, SM-DD and EXP, five days before the beginning of the anthesis. A series consists in two SM male inflorescence planted in single (SM-SD) and double density (SM-DD) respectively and three EXP male inflorescence sampled in the field. A total of 31 SM PAMI and 43 EXP PAMI were sampled and studied.

The anthesis stage was classified according of the flowers opening from the base to the top of the male inflorescence (Chiu et al. 1986) The anthesis starts with the opening of the first florets at the base of the spikelets. This gradual opening progresses upwards until all florets in the spikelets have opened in about 5-6 days. As soon as the anthesis stage started (stage M_1_), the anthesis stages (stages M_1_ to M_5_) were observed each day in the field until the end of the anthesis after the *EK* females laid the last eggs (when there are no longer any insects attracted to the inflorescence).

Following Ponnamma et al. (2006), the first emergences start within 10-13 days (M_10_-M_13_) after the first day of male inflorescence anthesis (MIA). Three days after the end of the anthesis (stage M_8_), the five PAMI of a series are cut at the base of the inflorescence peduncle. They are stored and transported individually in a bag from the field to the laboratory, mentioning the row and the palm numbers, the planting material, the date of collection, the density of plantation for the SM inflorescences.

Immediately after collection, all the spikelets for each PAMI of the series were cut one per one from the base to the top of the PAMI by using a pair of pruning shears (Bahco®, France). The useful length of each spikelet (useful part = part with flowers) was measured and recorded by using a stainless-steel ruler (Maped®, France) from the first flowers at the base of the peduncle to the base of the spine of the spikelet. The total number of spikelets, the total length of all spikelets and the average length of spikelets was calculated for each inflorescence.

All the spikelets of one individual inflorescence (PAMI) were then introduced in a labelled individual cage (600×300×300mm) for progeny emergence. Five sides of the cage are made of mosquito netting (mesh size approximatively 500 µm) and one side are made of plastic for easy observation (Entomo-Silex, Paimboeuf, France). The cages are lightweight, foldable and the zip closure allow to collect easily *EK* imagos. A thermo hygro buttom (Wimesure®, France) was hooked outside of one of the five cages to record temperature and relative humidity during the entire period of the *EK* emergence, to get local environmental conditions around the cages. To prevent invasion of predatory ants, the cages containing the spikelets are placed high up on box containing water.

The assessment of *EK* which have emerged during the last 24 hours from PAMI were carried out by sampling the weevils at 10:00 am using a 50 ml tube. Only the weevils belonging to the *Elaeidobius* genus were sampled, not the other flower-emerging insects.

The operation is repeated every 24 hours until four days without any more emergences and the spikelets are left in the cage few more days to confirm there will be no more emergence.

The weevils are stored in a tube indicating the references for each PAMI (row, palm number, planting material, and sampling date). When the collection is completed, the spikelets are brewed altogether to collect dead individuals at the bottom of the cage and to record them as daily emerging weevils.

Each tube is frozen (-20°C for 2-3 hrs.) to get motionless insects easy for sex determination and counting (Fig. 1) (Latip et al. 2019).

The sex ratio of progenies that emerged from the entire PAMI (= comparative ratio of females and males in a sexually reproductive species) was quantified for each inflorescence studied.

Five replicates for each series were studied per year, in January, April, July, September and November in 2020, 2021 and 2022 respectively making a total of 74 PAMI samples (31 SM and 43 EXP). However, in January and April 2020, the replicate was only composed of 2 SM and 2 EXP PAMI. Due to rat attacks, the replicate in April 2021 considered one more EXP PAMI.

Rainfall data was obtained from four manual gauges recording independently the daily rainfalls at four different locations in the estate where the study was conducted. A daily rainfall was obtained by averaging over the four devices, and the cumulated monthly rainfall was calculated based on this daily average.

### Statistical analysis and data visualization

Statistical analyses and data visualization were carried out with R software (version 4.5.1, R Core Team 2025). All graphics were produced using the ggplot2 R package (Wickham, 2016).

In order to analyze more finely the *E. kamerunicus* emergence kinetics, we calculated the fraction of emerged insects over time per inflorescence and per *EK* sex, as follows:

For each inflorescence (i), harvested on cultivar (c), and the *EK* sex (s), the cumulative emergence fraction F at day (j) was calculated as:

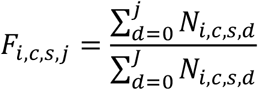

Where N_{i,c,s,d}_ is the number of insects of sex (s) that emerged from inflorescence (i) of cultivar (c) on day (d), and (J) is the last day of the emergence monitoring period. Thus, F_{i,c,s,j}_ represents the fraction of the total number of insects of sex (s) that had emerged by day (j).

We then modeled this cumulated fraction of emerged insects along time as a three-parameter logistic function:

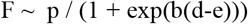

Where F is the cumulated fraction of emerged *EK*, p is the ceiling with a fixed value of 1, b is the slope and e is the lc50 parameter, d is the time from the start of emergence in days, and where b and e depends on the cultivar (c) and the *EK* sex (s) as fixed effects:

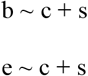

The model was fitted using the drm function from the drc R package (Ritz et al. 2015).

## Results

### Comparison of male inflorescence morphology between cultivars

Three morphological variables of the spikelets of the PAMI are presented (Fig. 2).

**Figure 2.**
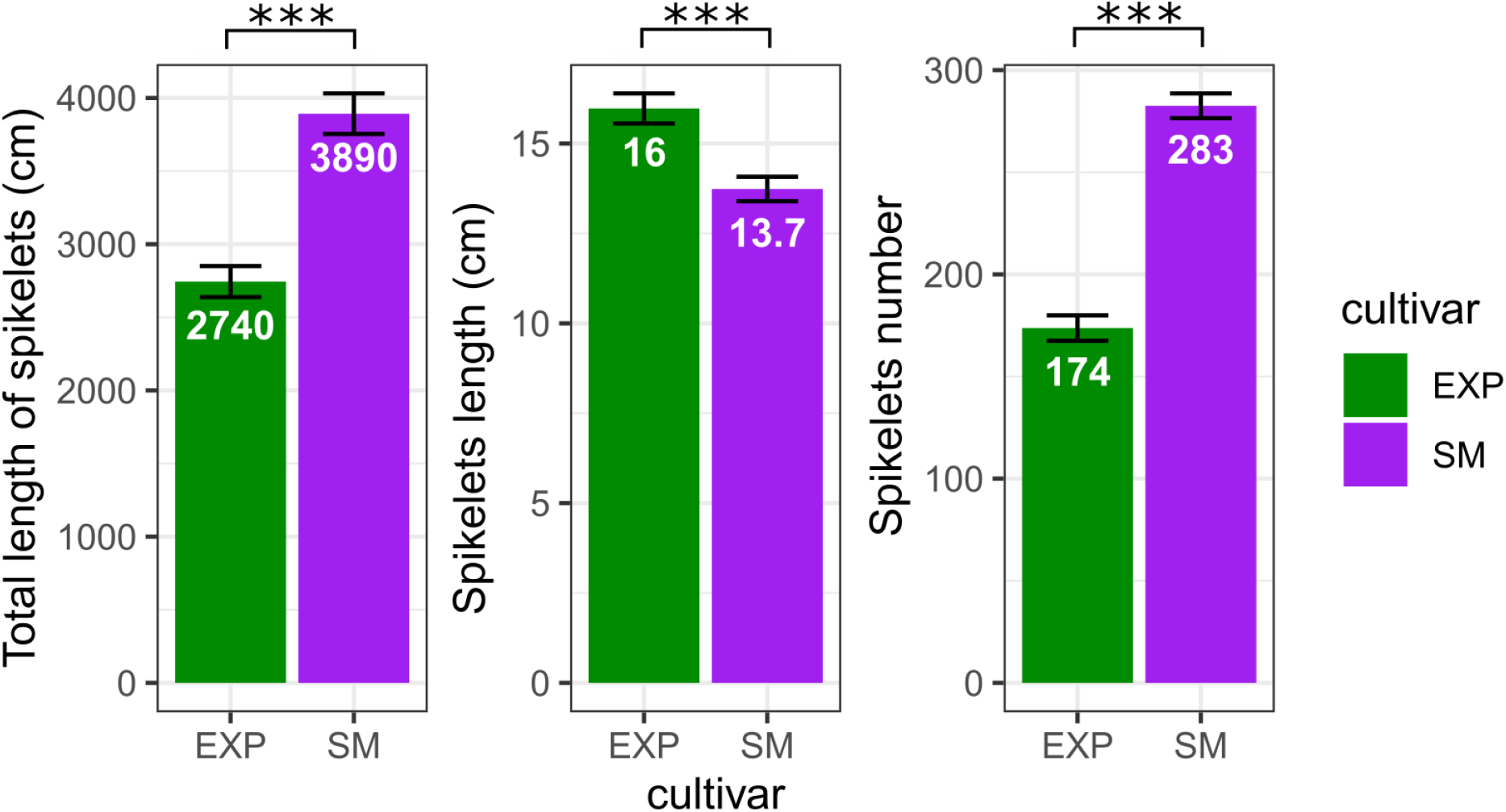
Differences in PAMI morphology measurements. Planting material = Cultivars (EXP: experimental cultivar, SM: super male). The data represent the distribution of mean values per inflorescence (n=43 for EXP and n=31 for SM). The figures in white represent the mean, the error bars represent the standard deviation and the stars represent the t-test p-value (***: p-value <0.001).

The average total spikelet length per inflorescence, measured on 43 EXP and 31 SM palms, was significantly greater in SM than in EXP (39 m vs. 27 m; *p =* 1.27 × 10⁻⁸) (Fig.2).

A similar pattern was observed for the average number of spikelets per inflorescence. SM palms produced significantly more spikelets per inflorescence than EXP palms, with averages of 283 and 174 spikelets, respectively (*p = 1.26 × 10⁻¹⁹*). Overall, these results indicate a significant effect of cultivar on both total spikelet length and spikelet number per inflorescence (*p <* 0.05; Fig. 2).

By contrast, although SM palms produced more and longer total spikelets per inflorescence, individual spikelets were significantly longer in the EXP cultivar than in the SM cultivar (16 cm vs. 14 cm; *p =* 8.92 × 10⁻⁵) (Fig. 2).

Analysis of total spikelet length and the number of spikelets per inflorescence over the three-year observation period showed that both traits increased between 2020 and 2022, indicating a significant effect of year (Fig. 3).

**Figure 3.**
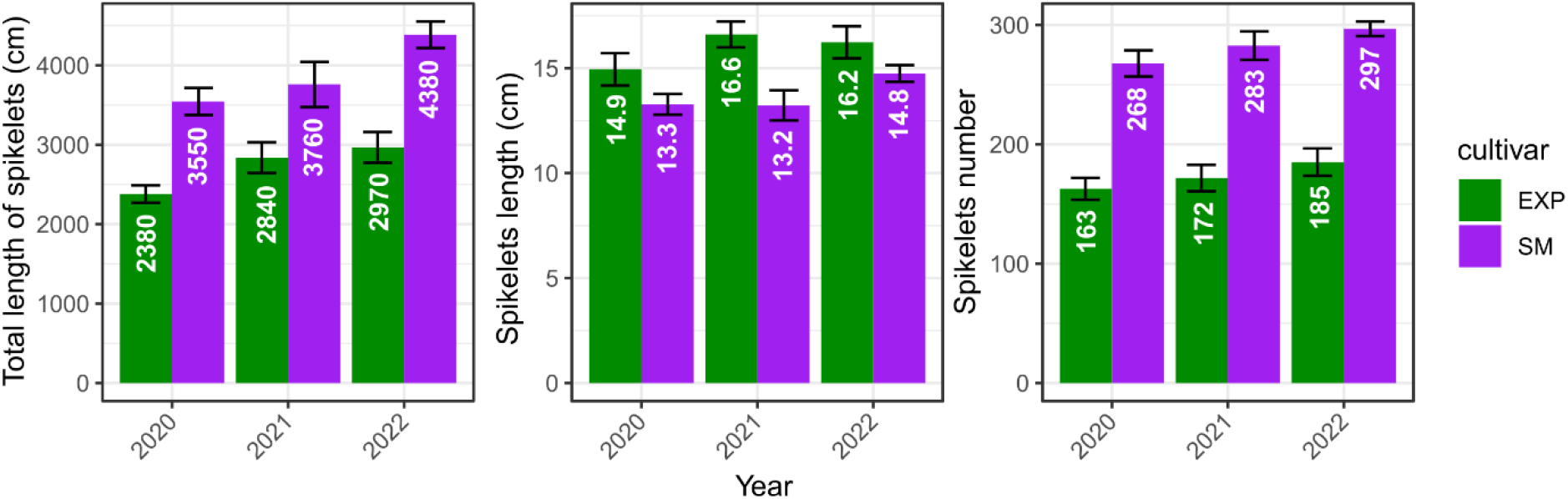
Differences in PAMI morphology measurements, broken down by year and type of cultivar. Planting material= Cultivars (EXP: Experimental cultivar, SM=supermale). The data represent the distribution of mean values per inflorescence (n=43 for EXP and n=31 for SM. The figures in white represent the mean, and the error bars represent the standard deviation. The measures were taken from 2020 to 2021 which corresponds to palms aged 6 to 8 years after planting.

The total spikelet length per inflorescence increased slightly over the three years for both cultivars. However, SM consistently exhibited a significantly greater total spikelet length than EXP. Mean values increased from 3550 to 4380 cm in SM and from 2380 to 2970 cm in EXP (Fig. 3).

In contrast, the mean length of individual spikelets was consistently greater in EXP than in SM throughout the study period. Spikelet length remained relatively stable across years, averaging 14.9–16.6 cm in EXP compared with 13.2–14.8 cm in SM, suggesting that the cultivar effect was more pronounced than the year effect for this trait (Fig. 3).

Similarly, the number of spikelets per inflorescence increased slightly over time in both cultivars, with SM consistently producing substantially more spikelets than EXP. The number of spikelets increased from 268 to 297 in SM, whereas EXP increased from 163 to 185. This difference parallels the greater total spikelet length observed in SM (Fig. 3).

### Comparison of *E. kamerunicus* emergence from male inflorescences between cultivars

Four variables were calculated from the emergence data for both cultivars and subsequently compared. The results are presented in Fig. 4.

**Figure 4.**
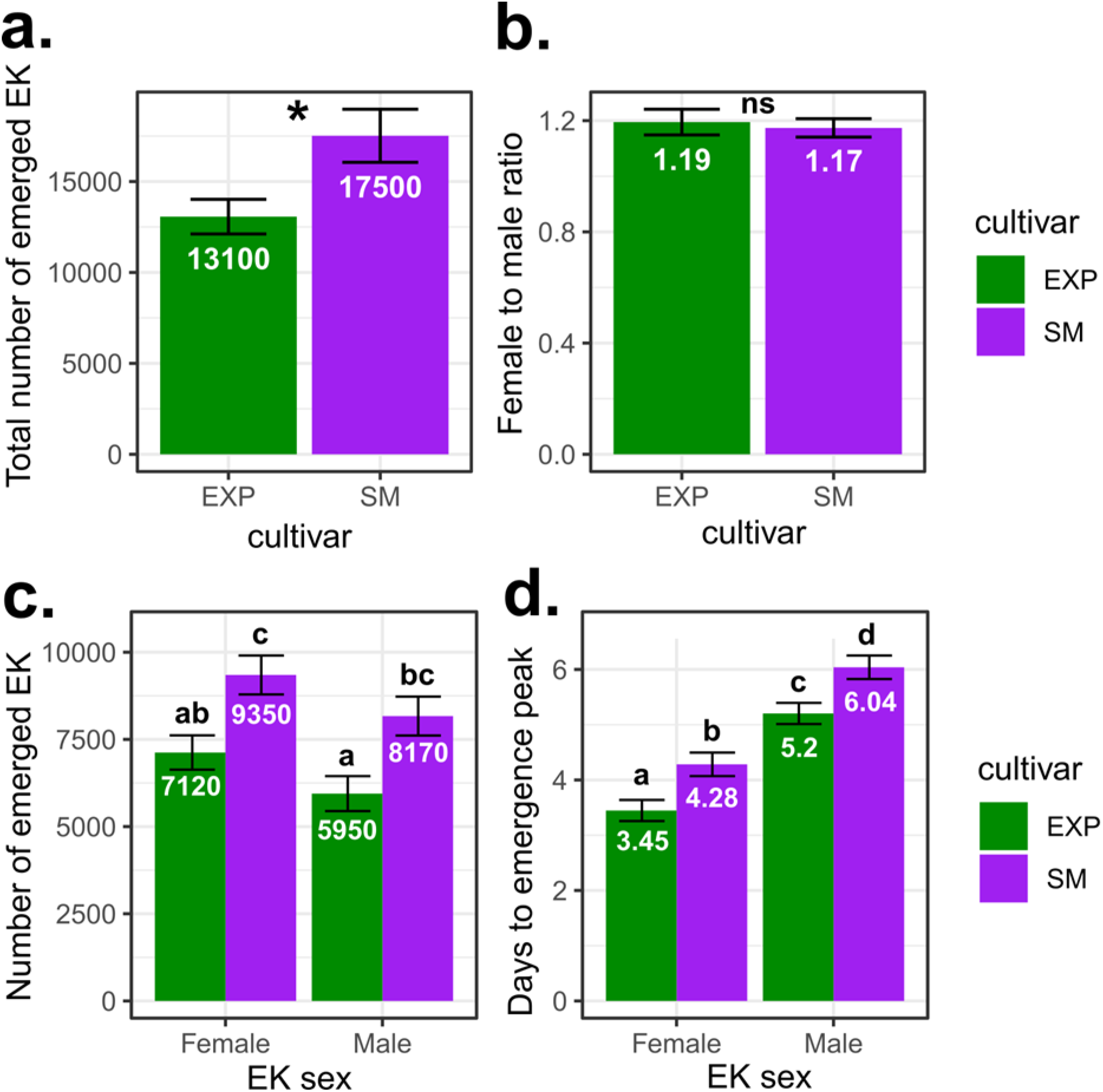
: Comparison of emergence characteristics between the two cultivars. EXP: experimental cultivar, SM=supermale cultivar. The data represent the distribution of mean values per inflorescence (n=43 for EXP and n=31 for SM). The figures in white represent the means (a. and b.) or estimated marginal means (c. and d.), and the error bars the standard deviation. Mean comparisons were performed either by t-test (a. and b.), or on estimated marginal means with EK sex and cultivar as fixed effect taking into account that the interaction wasn’t significant, using a post-hoc Tukey test with a confidence interval of 95% and alpha=0.05 (c. and d.). The data represent the distribution of mean values per inflorescence (n= 43 for EXP and n = 31 for SM).

The mean total number of *EK* adults emerging per inflorescence (both sexes combined) was significantly higher in the SM cultivar than in the EXP cultivar, with averages of 17,500 and 13,100 individuals per inflorescence, respectively (*p =* 0.013) (Fig. 4a). Among the 31 SM inflorescences examined, more than 30,000 *EK* adults emerged from three PAMI, between 20,000 and 30,000 from seven PAMI, between 10,000 and 20,000 from 15 PAMI, and fewer than 10,000 from six PAMI. In the EXP cultivar, among the 43 inflorescences examined, between 20,000 and 25,000 *EK* adults emerged from eight PAMI, between 10,000 and 20,000 from 17 PAMI, and fewer than 10,000 from 11 PAMI.

The sex ratio (female-to-male ratio) of emerging *EK* adults did not differ significantly between the two cultivars (*p =* 0.713). Both cultivars exhibited a female-biased sex ratio, with female-to-male ratios of 1.19 and 1.17 in EXP and SM, respectively (Fig. 4b).

The mean total number of emerging male *EK* adults was significantly lower than that of females in both materials. Mean male emergence was 5,950 and 8,170 individuals per inflorescence in EXP and SM, respectively, compared with 7,120 and 9,350 females (*p*= 0.015 for males; *p*=0.014 for females) (Fig.4c).

The time to peak emergence of *EK* adults differed significantly according to both sex and cultivar (Fig. 4d). Female adults reached peak emergence earlier than males in both cultivars. In the EXP cultivar, peak emergence occurred after an average of 3.45 days for females and 5.20 days for males. In the SM cultivar, the corresponding values were significantly longer, averaging 4.28 days for females and 6.04 days for males. Thus, *EK* female emerged earliest in the EXP cultivar, whereas *EK* male reached peak emergence latest in the SM cultivar.

### Dynamics of *EK* adult emergence

We observed two different temporal patterns of emergence between the *EK* sexes throughout the emergence period. The emergence period was longer for males than for females (Fig. 5a). Male *EK* emergence was spread over time, with individuals continuing to emerge until 20 days after the onset of emergence at the PAMI stage. In contrast, female emergence was concentrated within a shorter time frame, with most individuals emerging during the first 7 days. Emergence then declined from day 8 onward and reached very low levels by day 14.

**Figure 5.**
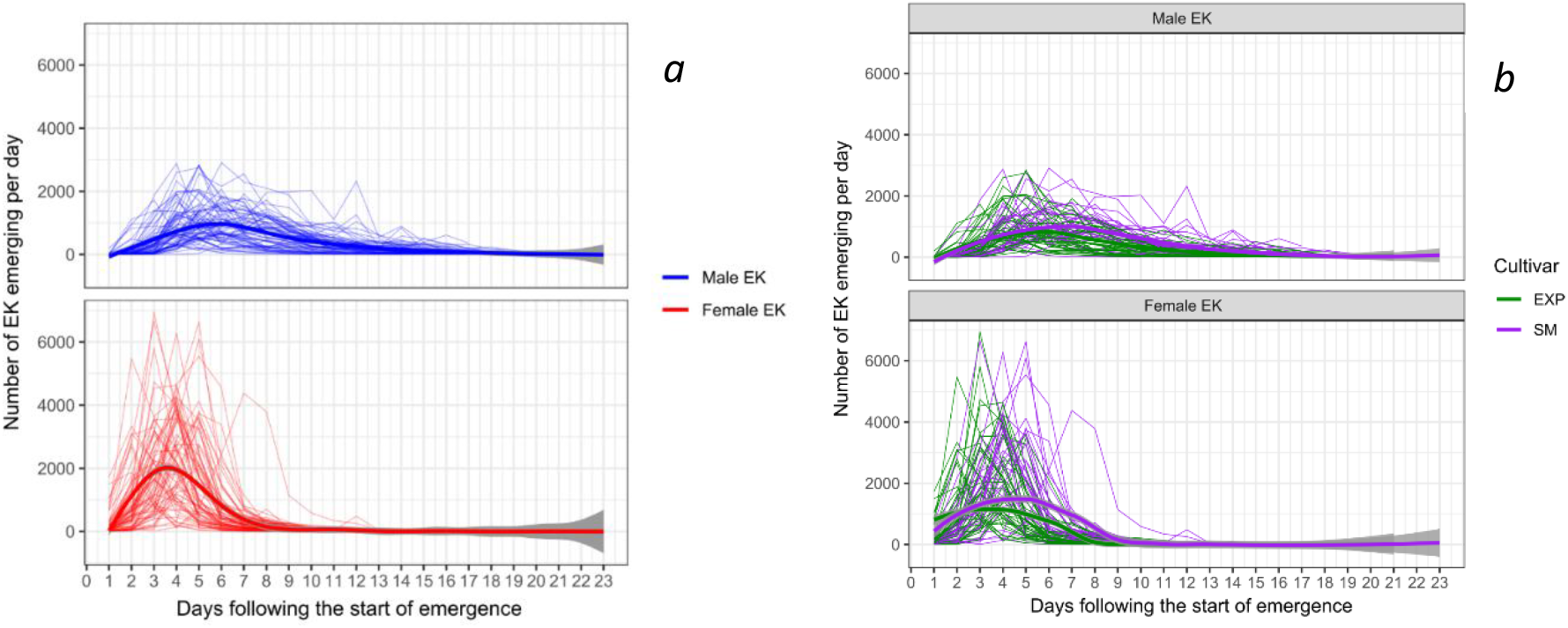
Dynamics of emergence over time (d_1_ = day with first emergences). Comparison of male (upper panel) and female (lower panel) E. kamerunicus, either regardless of cultivar (a) or according to cultivar (b). The thin lines represent the data for each individual inflorescence, and the thick lines represent a tendency line obtained by the local polynomial regression fitting as implemented in ggplot2 with the geom_smooth (method=”loess”) parameter including a confidence band (level=0.95) shown in grey.

Female emergence exhibited a marked peak 3-4 days after the start of emergence, with more than 6,000 individuals emerging during this period. In contrast, male emergence reached its maximum later, 5-6 days after the start of emergence, with a substantially lower peak of approximately 2,000-3,000 individuals (Fig. 4d and Fig. 5a). There seems to be no marked effect of the cultivar on the emergence kinetics (Fig. 5b).

In order to study more finely the differences in emergence kinetics between *EK* sex and between cultivars, we calculated for each inflorescence the cumulated fraction of emerged insects along time and we modelled this variable using a three-parameter logistic function in which the slope and lc50 parameters were allowed to vary according to *EK* sex and cultivar (Fig. 6). A significant difference of 2.65 days in lc50 (the time required to reach 50% cumulative emergence) was observed between males and females *EK* (*p* < 2 x10^-16^), indicating that female emergence peaked earlier than male emergence. In addition, the emergence curve was significantly steeper for females than for males (difference in slope = 0.61), suggesting a shorter emergence period for females (Fig. 6). The cultivar had a significant effect only on lc50 (*p* < 2e^-16^), with a difference of 0.74 days observed between EXP and SM, indicating that the emergence peak occurred slightly later in SM than in EXP palms.

**Figure 6.**
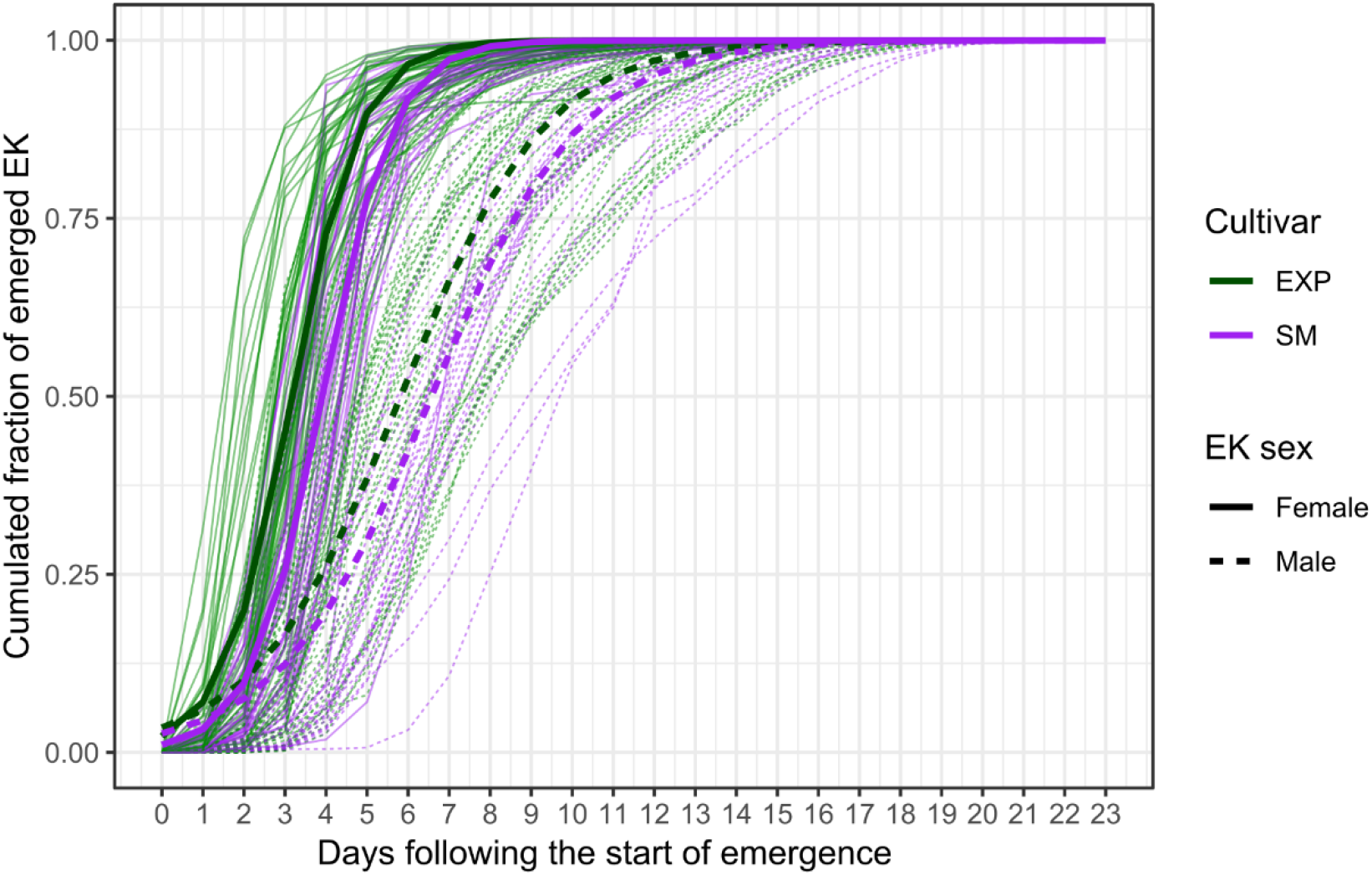
Emergence kinetics per cultivar and EK sex. The thin lines represent the cumulated fraction of emerged EK per inflorescence and per sex over time, in days from the start of emergence. Tendency lines (thick lines) were calculated using a three-parameter logistic function.

Overall, this analysis showed that *EK* sex had a major effect on the emergence dynamics, while the cultivar only had a minor effect, and the predicted emergence kinetics indicated that 95% of *EK* females emerged within 6-7 days compared with 11-12 days for 95% of *EK* males (Fig. 6).

Thus, analysis of the cumulated fraction of emerged weevils and its modelling using a logistic regression clearly confirmed that female and male emergence occurred at different times (Fig. 6), allowing a more accurate estimation of the temporal shift between the sexes.

### Factors associated with *EK* total emergences

We then studied whether variations were observed among months and years over the whole study period. The average number of *EK* emerging from January 2020 to December 2022 showed that *EK* emergence occurred throughout the five months of the study period, although its intensity varied over time (Fig. 7). A marked emergence peak was observed in late 2020 (approximately 22,000 individuals emerging on average in September), followed by a gradual decline until early 2022 (less than 15,000 individuals on average) and a subsequent increase during mid-2022 (more than 17,500 individuals emerging on average in April, July and September). Monthly rainfall also showed substantial temporal variation among months and years, with several periods of heavy rain occurring in late 2020 (420 mm in November), late 2021 (500 mm in November) and late 2022 (490 mm in November).

**Figure 7:**
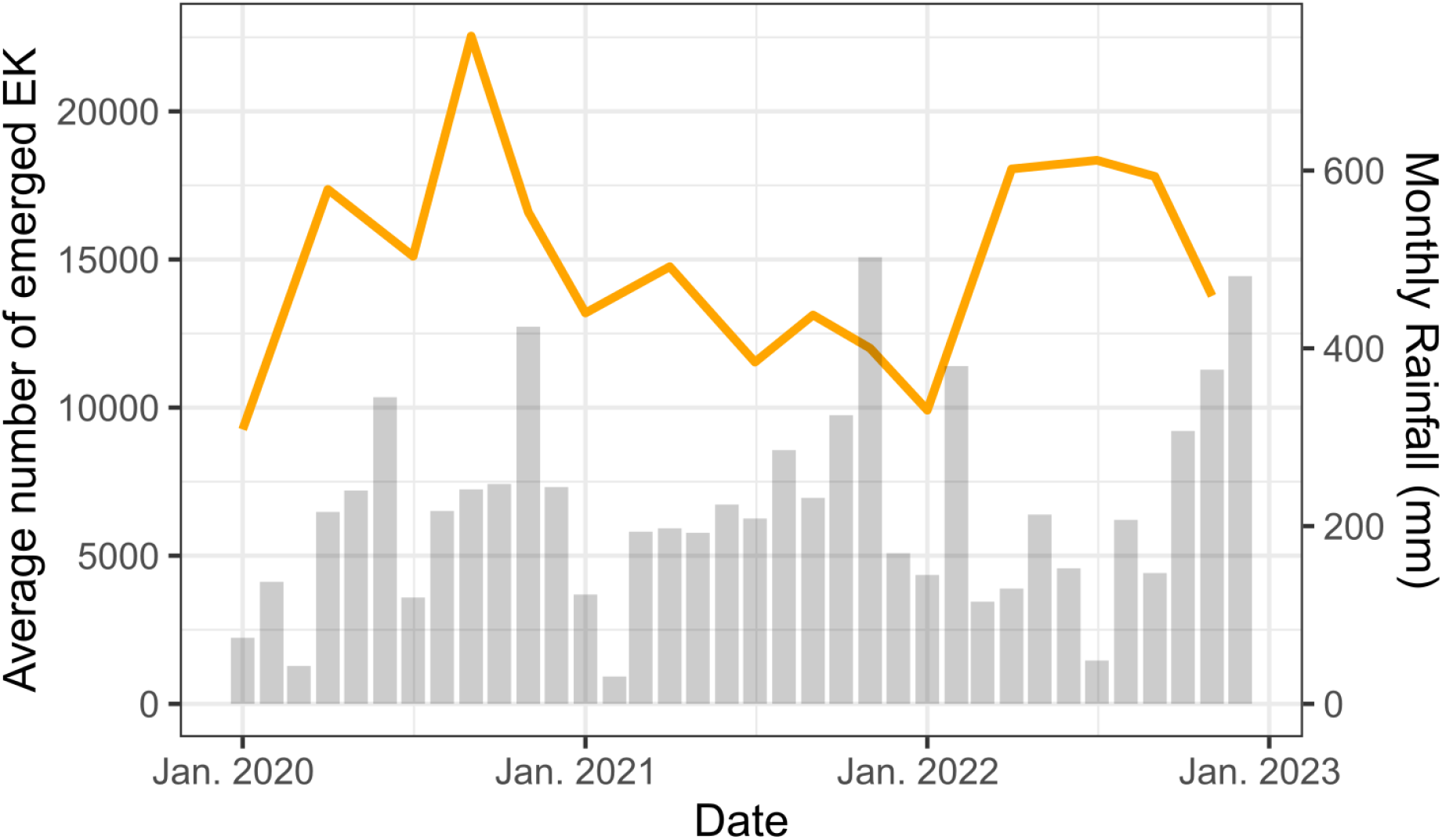
Temporal dynamics of EK emergence and rainfall from January 2020 to December 2022. The orange line shows the monthly average number of emerging EK individuals per inflorescence whereas the grey bars represent monthly rainfall (in mm).

No clear temporal relationship was observed between rainfall and *EK* emergence. Nevertheless, periods of maximum rainfall consistently coincided with lower *EK* emergence, while the highest emergence levels were recorded during periods of moderate rainfall.

Based on this result, we investigated whether *EK* populations could be influenced by rainfall occurring before male inflorescence anthesis. To this end, we tested the relationship between *EK* adult emergence and the cumulative rainfall recorded during the two-and three-month periods preceding *EK* emergence. In both cases, the correlation was negative (*r* = - 0.35), but this was not statistically significant for either the two-month (*p* = 0.21) or the three-month period (*p* = 0.20). We also observed no significant correlation between the total number of *EK* adults emerging per inflorescence with the cumulative rainfall recorded during the eight days preceding the collection of the inflorescence at the post-anthesis stage (Fig. 8).

**Figure 8.**
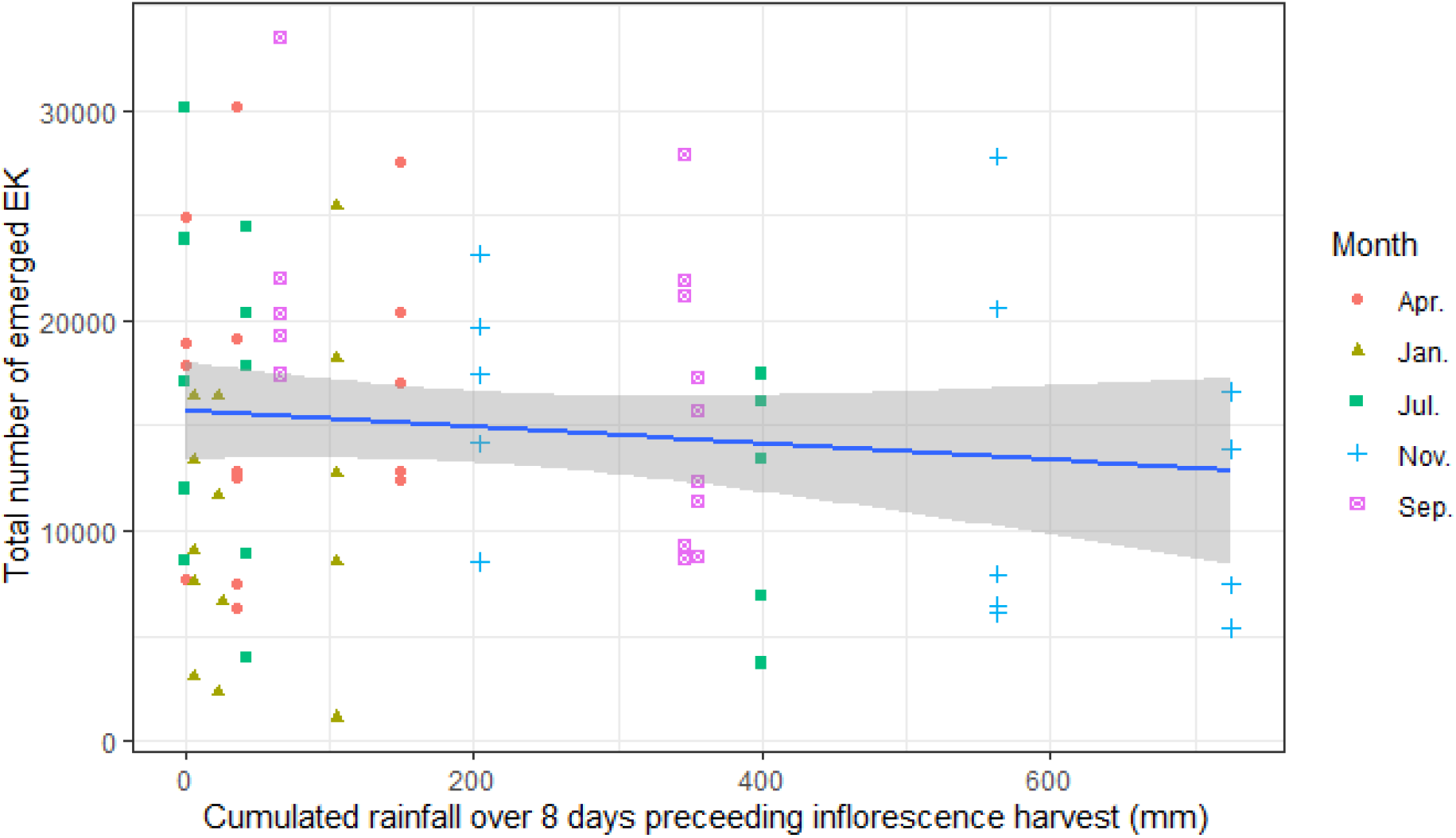
Total EK emerged and cumulated rainfall over 8 days preceding inflorescences harvest. The total number of emerged EK per male inflorescence is represented by points with shape and color depending on the month. The linear regression is presented as a blue line with the 95% confidence interval in grey as implemented by the geom_smooth (method= ‘lm’) function.

We assessed the relationship between total spikelet length and total number of emerging *EK* individuals (Fig. 9). A significant positive correlation was observed in the overall dataset as well as within each cultivar.

**Figure 9.**
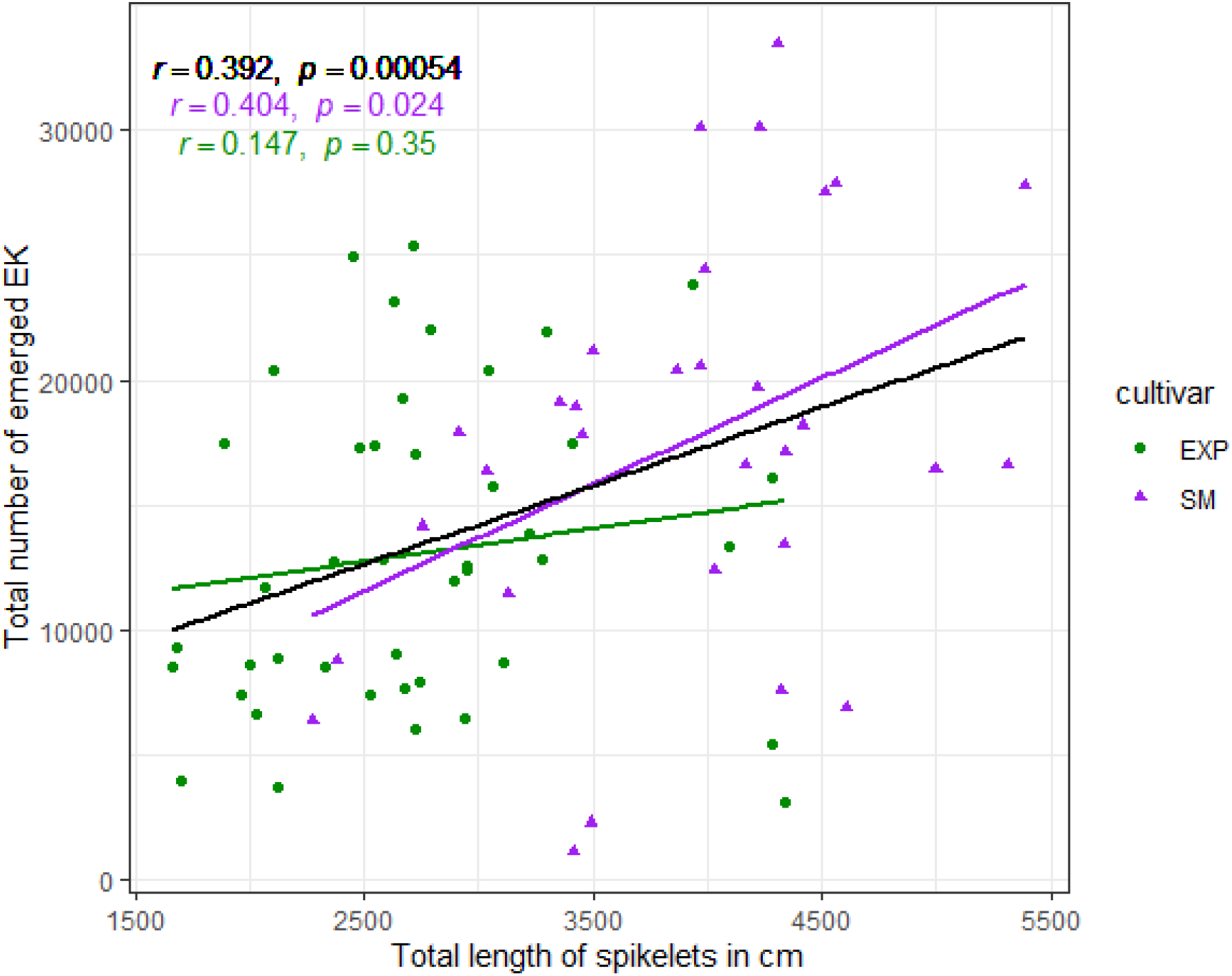
Correlations between total spikelet length and total number of emerging EK. The points correspond to the measures for each inflorescence (n= 43 for EXP and 31 for SM), the lines the regression lines with the Pearson correlation coefficients (r) and associated p-value (p, from t-test) indicated at the top left corner.

A positive relationship was observed in the overall dataset between total spikelet length and the total number of emerging *EK* (*r* = 0.392, *p* = 0.00054). This correlation was observed in both SM and EXP, confirming that total spikelet length influences the number of emergences.

When the two cultivars were analyzed separately, this relationship remained significant for the SM cultivar (*r* = 0.404, *p* = 0.024), whereas no significant correlation was detected for the EXP cultivar (*r* = 0.147, *p* = 0.35). The steeper regression slope for SM suggests that increase in spikelet length are associated with a greater increase in the number of emerging *EK* than in EXP. In contrast, the weak and non-significant relationship observed in EXP indicates that spikelet length is a poor predictor of emergence in this cultivar. A positive correlation was also found between *EK* emergence and the number of spikelets (*r* =0,367, *p* = 0,00128).

## Discussion

This study investigates and compares the number of successfully emerged adult *EK* from PAMI in North Sumatra between two types of cultivars (EXP and SM). It also evaluates the effects of spikelet length, spikelet number, and meteorological conditions on progeny emergence.

The mean number of spikelets per inflorescence was 174 for EXP and 283 for SM, corresponding to palms aged between 6 and 8 years after planting, respectively. These values are consistent with those reported by Beaudoin-Ollivier et al. (2026). SM male inflorescences exhibited a greater number of spikelets per inflorescence than that of EXP palms. For EXP, the mean value is comparable to the average of 170 spikelets reported by Chiu et al. (1986) whereas for SM it falls within the range of 100 to 300 finger-like spikelets described by Jacquemard et al. (2011). The number of spikelets per inflorescence exhibits considerable variation among palms. However, Beirnaert (1935) reported that variation among inflorescences within an individual palm is relatively limited. In the present study, both total spikelet length and spikelet number were significantly affected by the year (2020, 2021 and 2022) which is confounded with palm age. SM consistently exhibited a significantly greater total spikelet length than EXP indicate that SM developed a more extensive inflorescence architecture. Corley & Gray (1976) showed that the average number of spikelets increases with palm age, from approximatively 60 in 3-year-old palms to about 150 in 10-year-old palms. Similarly, Rahardjo et al. (2018) and Tandon et al. (2001) reported that older oil palms generally produce a greater number of spikelets. These observations suggest that *EK* abundance may increase with palm age. The number of spikelets per inflorescence differed only marginally between 5- and 13-year-old palms, with a slight increase observed in older palms (Siswanto & Soetopo 2020).

The mean total spikelet length was 27 m for EXP and 39 m for SM. These values are substantially higher than those reported by Beaudoin-Ollivier et al. (2026). This discrepancy may be explained by differences in spikelet morphology, as Beaudoin-Ollivier et al. (2026) included abnormal spikelets containing sections devoid of flowers, which likely reduced the total flowering spikelet length.

The mean length of individual spikelets in PAMI was 16 cm and 14 cm for EXP and SM respectively. These values are within the range reported by Corley & Tinker (2016), who described spikelet lengths of 10-20 cm and by Janick & Paul (2008), who reported values ranging from 10 to 30 cm. The mean number of emerged *EK* per unit of total spikelet length was 4,52 individuals cm^-1^ for SM and 4,93 individuals cm^-1^ for EXP with no significant difference observed between the two cultivars. Under laboratory conditions Ponnamma et al. (2006) reported a strong positive correlation between the number of emerged weevils and spikelet length (*r* = 0.4315). The authors suggested that longer spikelets may provide greater resources for pollinating weevils, including increased food availability and more suitable breeding sites, compared with shorter spikelets.

Morphological measurements of spikelets revealed contrasting inflorescence architectures between the two cultivars. SM exhibited a strategy based on producing greater number of spikelets, resulting in a higher cumulative spikelet length per inflorescence, whereas EXP produced fewer but longer spikelets, leading to a lower total spikelet length despite the greater average length of individual spikelets. These differences suggest distinct developmental strategies, with SM favoring spikelet proliferation and EXP favoring spikelet elongation. Interestingly, despite a slight increase in all three traits from 2020 to 2022, the differences between cultivars remained remarkably stable over time. This pattern suggests that genetic differences between cultivars may have a greater effect on inflorescence morphology than year-to-year environmental variation under the studied conditions.

### Comparison of emergence characteristics between cultivars

The originality of our study lies, first, in the comparison of the number of successfully emerged *EK* adults from PAMI between two different cultivars (EXP and SM) and second, in the analysis of the emergence potential of the entire PAMI progeny.

The total number of emergences recorded across the entire PAMI was significantly higher in SM palms (17,500) than in EXP palms (13,100). Compared with previous studies, the mean number of weevil emergences observed under our experimental conditions was substantially higher than that reported by others authors (Dhileepan 1994; Ponnamma et al. 2006; Mohamad et al. 2022, 2023). However, these results should be interpreted in light of the morphological differences between the two cultivars. SM palms produced more spikelets (283 on average), but these were shorter (14 cm), whereas EXP palms produced fewer spikelets (174 on average), although they were longer (16.0 cm). Rahmadani (2015) and Kurniawan (2010) states that the sufficient availability of a number of spikelets determine the abundance of the emerging population of *EK*.

Nevertheless, all the studies reviewed estimated weevil emergence from only a limited number of spikelets per inflorescence, typically three spikelets sampled at each inflorescence level. In our study, this means an average rate of emergence of 75 ± 40 *EK* adults per spikelet for EXP palms and 62 ± 28 *EK* adults per spikelet for SM palms. Ming & Bong (2017) reported a mean emergence of 43 *EK* adults per untreated PAMI spikelet. Similarly, Mohamad et al. 2023 and Saravanan et al. (2023) reported emergence rates ranging from 36 to 47 and from 35 to 87.3 *EK* adults per spikelet, respectively. In Malaysia, Syed (1982) reported a mean of 63.6 *EK* adults emerging per PAMI spikelet, whereas in Cameroon, the corresponding values were 50.7 *EK* adults and 67.3 adults of other *Elaeidobius* species. Furthermore, emergence per spikelet has been reported to range from 20 to 90 adults, with fewer than 30 adults recorded in two plantations and more than 160 adults per spikelet in Sabah (Syed et al. 1982). Likewise, Ponnamma et al. (2006) reported emergence values ranging from 25.86 to 134,02 per spikelet. These findings suggest that the cultivars used in our study have the potential to produce a higher number of *EK* individuals per spikelet than previously reported. This high emergence rate may also reflect the good health and reproductive performance of the *EK* population during the study period (2020 to 2022) but also more flowers occupied by *EK* immature stages due to a low density of male inflorescences available.

With mean emergence values of 17,500 *EK* adults per SM male inflorescence and 13,100 adults per EXP male inflorescence, our results suggest that both cultivars are able of supporting high levels of pollinator abundance and, consequently, favorable fruit set. According to Donough et al. (1996), a high oil palm fruit set (i.e., above 75%) requires a minimum population of approximately 20,000 *EK* individuals per hectare. More recently, Mohamad et al. (2021) showed that a population of 14,000 weevils per hectare was sufficient to achieve a fruit set of about 65%. However, the threshold appears to vary among production environments. For example, Wahid & Kamarudin (1997) reported that fruit set exceeding 60% could be obtained with as few as 4,711 weevils per hectare, whereas Dhileepan (1994) estimated that approximately 7,000 weevils per hectare were required under Indian conditions.

### Role of rainfall over an 8-day, 2- month and 3-month period before the emergence progenies record

September to November corresponds to the rainy season in the study area. Overall, the mean number of *EK* progeny emerging was not markedly affected by cumulative rainfall during the 8-day, 2-month, or 3-months periods preceding the harvest of the male inflorescences, despite the species generally being considered to prefer relatively dry conditions.

Previous studies have reported contrasting effects of rainfall on *EK* populations. Several authors suggested that the wet season and a favourable rainfall distribution promote population build-up (Hutauruk et al. 1984; Wahid et al. 1987; Syed & Saleh 1987; Dhileepan 1994; Nurul Fatihah et al. 2018, 2019). However, adult weevils are known to be less active during wet weather. Despite this reduced activity, *EK* was selected for introduction into Asia because its populations were less affected by the rainy season in Cameroon than those of other pollinating weevil species (Syed 1982). In the present study, the lowest mean progeny emergence was observed in January, although cumulative rainfall during the 8-days preceding the harvest of the PAMI varied widely (0-400 mm), suggesting that short-term rainfall alone does not explain this seasonal decline. Syed & Saleh (1987) reported that *EK* populations are negatively affected by several factors, including drought.

Dhileepan (1994) further observed that, although *EK* populations declined during the dry season, pollination efficiency was not reduced. The same author also found no correlation between total rainfall and progeny emergence, whereas the number of rainy days was positively correlated with progeny emergence. More recently, Saravanan et al. (2023) reported highly significant negative correlations between *EK* progeny abundance and minimum, maximum, and mean temperatures, as well as rainfall.

Several factors may influence weevil emergence, including palm age, the period of observation, the proportion of inflorescences available, and the distribution of insects according to the position of the spikelet within the inflorescence (Siswanto & Soetopo 2020). Syed (1982) further reported that *EK* progeny is distributed throughout all regions of the male inflorescence and along the entire length of each spikelet.

The number of successfully emerging progeny may also be affected by biotic factors such as predation by rats, nematodes, mites and other natural enemies (Luqman et al. 2017). In addition, the application of insecticides directly to male inflorescences can adversely affect the activity, survival, and reproductive success of pollinating weevils (Prasetyo & Susanto 2019, Mohamad et al 2021). Under laboratory conditions, insecticide treatments significantly reduced the number of newly emerged weevils per inflorescence spikelet. Similarly, Ming & Bong (2017) reported that insecticide applications for bunch moth control in young oil palm plantations had significant detrimental effects on both adult weevil survival and, more markedly, the emergence of new weevils from PAMI.

Although our results did not reveal a clear relationship between rainfall and *EK* emergence, previous studies have suggested that seasonal rainfall may influence the species’ reproductive performance. Dhileepan (1994) reported that the mean number of *EK* progeny produced per PAMI spikelet varied markedly among seasons., ranging 4.2 to 46.6 adults per spikelet. Likewise, Nurul Fatihah et al. (2019) found that the rainy season provides favorable conditions for the development to *EK*. Similar observations were also reported by Hutauruk et al. (1984); Syed & Saleh (1987); Wahid et al. (1987) and Dhileepan (1994). It’s also known from observations in the field that intense drought can result in PAMI drying, with an important negative impact on the developing *EK* larvae.

### Sex ratio of newly emerged progenies

The mean number of newly emerged males differed significantly between the two cultivars, being lower in EXP palms (5,950 individuals per male inflorescence) than in SM palms (8,170 individuals per male inflorescence). A similar pattern was observed for females, with significantly fewer individuals emerging from EXP palms (7,120 individuals per male inflorescence) than from SM palms (9,350 individuals per male inflorescence). In both cultivars, female emergence exceeded male emergence, resulting in a female-biased sex ratio. This finding is consistent with previous studies (Saravanan et al. 2023; Yue et al. 2015).

Despite differences in the total numbers of emerging adults, the sex ratio of newly emerged progeny did not differ significantly between the two cultivars, averaging 1.19 females:1 male in EXP palms and 1.17 females:1 male in SM palms. These values are within the range reported by Ponnamma et al. (2006) who observed female-to-male ratios ranging from 1.13:1 to 1.94:1, although they are lower than those reported by Mohamad et al. (2023), where the ratio ranged from 2.33:1 to 4.46:1.

Several hypotheses have been proposed to explain the predominance of females in *EK* populations. A female-biased sex ratio may increase reproductive success when the production of females contributes more strongly to population growth, particularly under conditions where a limited number of males can fertilize multiple females (Charnov 1982). More recently, Rodri’guez-Mum̃oz et al. (2019) proposed that female-biased sex ratios in wild insect populations may also be associated with slower female senescence. In several insect species, intracellular bacteria can profoundly alter sex ratios by biasing offspring production towards females (Werren et al. 2008). Because females are the reproductive component of the population, the sex ratio is considered an important demographic parameter that can influence reproductive success, population growth, and long-term population dynamics.

### Dynamics of newly emerged progenies from PAMI

Our results showed that adult *EK* emergences occurred throughout the year, with emergence events recorded in January, April, July, September and November over three consecutive years (2020-2022). Adult *EK* emergences was distributed over an average period of 17 days for both males and females. The duration of emergence from individual inflorescences was similar between EXP and SM PAMI, ranging from 10 to 23 days. These observations are consistent with Ponnamma et al. (2006) who reported that weevils began emerging within 10-13 days under laboratory conditions.

Despite a fairly balanced sex ratio, the emergence dynamics between males and females are clearly different. The findings of the emergence distribution clearly show that females begin to emerge before males and in a larger number, irrespective of cultivar, suggesting protogyny in adult emergence (i.e., females become available before males). Then, the emergences contained a mixture of males and females. The emergence of males spread over a longer period of time (16 days) than that of females, which occur in clusters (6 days). From the day 12 after the first emergence started, there are still males emerging whereas no more female emerged. Is there an emergence strategy?

In addition, emergence was more spread out over time in SM cultivar with lc50 delayed by 0.8 day compared with the EXP cultivar. This delay may reflect differences in inflorescence characteristics or in the developmental conditions provided by the two cultivars.

Females reached their emergence peak significantly earlier than males across both cultivars. Compared with EXP, the SM cultivar delayed the timing of peak adult emergence in both sexes. The consistent difference between females and males, together with the cultivar effect, suggests that both sex and host plant genotype influence the developmental timing of *E. kamerunicus*. Such differences in emergence dynamics may affect pollinator availability and synchrony with anthesis, potentially influencing pollination efficiency.

Female emergence before male emergence (protogyny, used by Buck 2001) occurs in approximately 36% of the 192 species for which data are available (Teder et al. 2021). This strategy is wide-spread in bagworms (Lepidoptera: Psychidae) (Rhainds et al. 2009). Sexual differences in development time are imperfectly understood. An abundance of experimental data exists on the length of male and female pre-imaginal (i.e. pre-adult) development, largely on insects of economic importance (Honek 1997).

The observed differences in emergence timing between sexes may be explained by a combination of ecological and evolutionary factors, including (i) reproductive strategy and mate availability, (ii) resource competition and oviposition site selection, and (iii) environmental and developmental constraints.

First, earlier emergence of females may represent an adaptative strategy to optimize reproductive success. Adult female insects generally require a period of feeding to complete egg maturation before reproduction (Jarosik & Honek 2007). Consequently, females emerging earlier can reach reproductive maturity before the arrival of males, allowing immediate mating once males become available. Delayed male emergence may ensure that males encounter already receptive females while avoiding premature emergence when reproductive opportunities are limited. Such temporal synchronization may maximize fertilization success, enhance population reproductive output, and potentially reduce the risk of inbreeding (Rhainds & Gries 1995). A lower emergence success of males compared with females has also been reported in *Curculio elephas* (Coleoptera: Curculionnidae) (Menu 1993).

Second, because *EK* larvae develop within oil palm tissues (spikelets), early female emergence may provide a competitive advantage by allowing females to access and secure suitable oviposition sites before resources competition increases (Degen et al. 2015).

Once females have occupied resource-rich sites, later-emerging male may locate them through pheromone cues or visual/olfactory signals. Therefore, earlier female emergence may enhance access to high-quality reproductive resources and improve offspring production.

Third, differences in developmental rates between sexes may also contribute to asynchronous emergence. Females may complete larval and pupal development more rapidly than males, potentially due to differences in developmental pathways, including a possible reduction in the number of larval instars. In contrast, males generally require a longer maturation period because of their larger body size (Latip et al. 2019). A positive relationship between development time and body size has been reported by Roff (1980).

In holometabolous insects such as *EK*, the pupal stage may contribute to slower male development compared with females. This difference could result from the higher energetic investment required for preimaginal development of male reproductive organs, making male gonad development more costly than that of females (Blanckenhorn et al. 2007). Such increased developmental costs may contribute to the lower rate of male maturation. Microclimate effects (temperature, humidity) can also differentially affect male vs. female development rates. In the last case, early female emergence, followed by male emergence helps regulate the mating dynamics.

The main hypothesis according to Buck (2001), in necrophagous Diptera, is that bimodal emergence could have the advantage of maintaining survival chances in the event that emerging specimens face unfavorable conditions.

Otherwise, it is difficult to justify precisely this delay between the emergence of females and males. Several hypotheses other than those of Buck (2001) are possible. One of them is based on the fact that the variability in the timing of emergence is genetically controlled in the genus *Trichogramma* (Pompanon et al. 1995).

Finally, the temporal difference in emergence between females and males could constitute a mechanism contributing to reducing the risk of inbreeding in some insects. In particular, asynchronous emergence of males and females, combined with post-emergence dispersal, may decrease the likelihood of mating between related individuals originating from the same breeding site. In the parasitoid *Aphidius matricariae*, emergence timing and adult dispersal have been associated with a reduction in opportunities for sibling mating (Bourdais & Hance, 2019).

### Number of newly emerged *EK* at PAMI and total spikelet length

When comparing EXP and SM PAMI, the total length of spikelets for one inflorescence affects the number of emerging *EK*. Dhileepan (1994) indicated emergence of *EK* increased with palm age due to the increase of spikelet length and numbers of spikelets.

## Conclusion

Our study reveals that by increasing floral resources, nesting habitat availability by using SM cultivar, the progenies emergence of *EK* increase. As the total length of spikelets and the number of spikelets per inflorescence are higher in SM cultivar, we may assume that SM may be useful to ensure a sufficient natural pollination particularly in young plantations. Demonstration was done that the number of progenies that successfully emerge from the SM post-anthesis male inflorescence is considerably higher compared to the EXP PAMI. To maintain oil palm yield production, SM cultivar is then recommended to sustain the *EK* population.

The method considering the emergence of the entire post-anthesis male inflorescence allowed to estimate the general health of *EK* population and to predict *EK* emergence kinetics.

Finally, our results revealed the strategy of *EK* females emerging before males. Protogyny is rare but not exceptional in insects. Its adaptative significance is difficult to establish because of the limited knowledge of the mating behavior of *EK*. Protogyny remains an emerging research topic, making it an active area of investigation in evolutionary ecology and insect development biology.

The sex ratio of newly emerged progenies is female-biased as recorded for the *EK* natural population trapped at the vicinity of male inflorescence in anthesis. A female-biased sex ratio suggests that specific ecological or evolutionary pressures are acting to modify the expected sex-ratio equilibrium.

Future studies must consider the communities of other insects that are emerging from entire post-anthesis male inflorescence as they were not taken into consideration in this present study.

## Acknowledgments

The authors would like to extend their gratitude to the executive director of PT SOCFINDO (SOCIETE FINANCIERE DES CAOUTCHOUCS INDONESIE), H.O. Williams for permission to publish this work. The authors wish to thank Centre international de La Recherche pour le Développement (CIRAD), PT SOCFINDO and PALMELIT SAS for their financial support. We would also like to thank the woman staff of the pollen laboratory of SOCFINDO SEED PRODUCTION AND LABORATORIES (SSPL) for their technical support as they were daily in charge to collect the weevils from the emerging cages, as well as the management of the estate and the tireless workers who assisted us in this study.

## Author contribution

LBO conceived protocol and research. DA, IS, LBO, YD, HR and CM conducted the experiments and collected data for daily emergence. FJ performed the statistical analysis. LBO and FJ wrote the original manuscript. YD and CM red and approved the manuscript.

## Generative AI disclosure statement

No generative AI in any way was used to produce this article.

